# Spatial mapping reveals unique cellular interactions and enhanced tertiary lymphoid structures in responders to anti-PD-1 therapy in mucosal head and neck cancers

**DOI:** 10.1101/2024.04.18.590189

**Authors:** A.L. Ferguson, T. Beddow, E. Patrick, E. Willie, M.S. Elliott, T.H. Low, J. Wykes, M.H. Hui, C.E. Palme, M. Boyer, J.R. Clark, J.H. Lee, U. Palendira, R. Gupta

**Author notes:** authors have contributed equally (A.L. Ferguson, T. Beddow). authors have contributed equally (J.H. Lee, U. Palendira, R. Gupta).

## Abstract

Survival in recurrent/metastatic head and neck mucosal squamous cell carcinoma (HNmSCC) remains poor. Anti-programmed death (PD)-1 therapies have demonstrated improved survival with lower toxicity when compared to standard chemotherapy. However, response to anti-PD-1 therapy remains modest, at 13-17%.

We evaluated the tumor microenvironment (TME) using Imaging Mass Cytometry (IMC) on 27 tumor specimens from 24 advanced HNmSCC patients prior to receiving anti-PD-1 based treatment. We show significantly increased central memory T cells and B cells in responders (n=8) when compared to non-responders (n=16). Spatial mapping identified interactions between phenotypically distinct malignant squamous cells with CD8+ T cells, CD4+ Tregs and endothelial cells in responders, and avoidance of these cells in non-responders. Importantly, regional analysis shows responders have more abundant tertiary lymphoid structures (TLS), with TLS proportion >20% also associated with longer progression free survival. Together these findings define the immune landscape associated with response to anti-PD-1 treatment in HNmSCCs.

---

The prognosis for recurrent/metastatic head and neck mucosal squamous cell carcinoma (HNmSCC) remains poor, with a median survival of less than 12 months. Systemic therapy is indicated in this setting, with the choice of regimen guided by clinical factors such as comorbidities, performance status, prior therapy, HPV status and programmed cell death ligand-1 (PD-L1) combined proportion score (CPS).

The anti-programmed cell death 1 (PD-1) agent pembrolizumab, either alone or in combination with chemotherapy, has demonstrated improvement in overall survival when compared to chemotherapy plus cetuximab (EGF-receptor targeted therapy) in untreated recurrent/metastatic HNmSCC patients in the Phase 3 KEYNOTE-048 (NCT02358031) clinical trial. However, Response Evaluation Criterial in Solid Tumours (RECIST) 1.1 response to pembrolizumab alone remains modest despite therapeutic stratification using PD-L1 CPS, with only 17% of the total population and 23% of patients with PD-L1 CPS of 20 or more responding to treatment (1). Despite the modest response rate, this was associated with durability of response, with the 24-month overall survival being 27% with pembrolizumab versus 19.1% for chemotherapy plus cetuximab in the total population(2). Another anti-PD1 therapy nivolumab has also demonstrated activity in the second-line setting with improvement in survival and lower toxicity compared to investigator choice second-line systemic therapy in the Phase 3 CheckMate-141 (NCT02105636) clinical trial (3).

In addition to PD-L1 CPS, biomarkers indicative of tumour inflammation such as tumour infiltrating lymphocytes and T-cell activation and IFN-γ gene expression signatures have been investigated (4, 5). A recent study explored the role of tumor mutation burden (TMB), T-cell-inflamed gene expression profile (GEP) and PD-L1 CPS in 257 patients receiving pembrolizumab as part of the KEYNOTE-012 and KEYNOTE-055 clinical trial(6). All three biomarkers were independently predictive of RECIST response, with the highest response observed in patients with high TMB plus high T-cell-inflamed GEP or PD-L1 CPS. Interestingly, this improved response did not correlate with improved progression free survival (PFS), with a trend for longer PFS observed in patients with high TMB only but not for high T-cell-inflamed GEP and PD-L1 CPS. Evaluation of the tumor microenvironment (TME) and the immune milieu of HNmSCC will complement biomarker investigations for predictors of response to immune check point inhibitors (ICI).

Currently, a detailed cellular map of the HNmSCC TME is not available in literature. Imaging mass cytometry (IMC) simultaneously allows high dimensional quantitative and spatial analyses of tumor cells and the TME using up to 40 antibodies. In this study we compared the TME of tumors collected prior to anti-PD-1 therapy from responding and non-responding patients with advanced/metastatic HNmSCC using IMC. We demonstrate significant differences in the proportions, densities and interactions of cells amongst responders versus non-responders, generating evidence to inform treatment strategies and to potentially overcome resistance mechanisms.

## Results

### Clinicopathological Characteristics

This study included 27 FFPE tumor specimens from 24 HNmSCC patients collected prior to treatment with anti-PD-1 therapy (n=3 patients had both primary and metastatic (LN) specimens). The median age was 56, 16/24 (67%) were male and location of primary included oral (n=11, 46%), oropharyngeal (n=10, 42%) and larynx (n=3, 12%). Of the ten patients with oropharyngeal SCC, five were p16 positive (Table 1).

**Table 1:** Baseline characteristics.

| Characteristics | Total<br>(n = 24) | Responders<br>(n = 8) | Non-responders<br>(n = 16) |
| --- | --- | --- | --- |
| Age | 58 (25–78) | 60 (54–73) | 58 (25–78) |
| Sex |  |  |  |
| Male | 16 (67%) | 5 (63%) | 11 (69%) |
| Female | 8 (33%) | 3 (37%) | 5 (31%) |
| Primary tumour location |  |  |  |
| Oral | 11 (46%) | 2 (25%) | 9 (56%) |
| Oropharyngeal | 10 (42%) | 5 (63%) | 5 (31%) |
| Laryngeal | 3 (12%) | 1 (12%) | 2 (13%) |
| p16 status |  |  |  |
| Positive | 5 (21%) | 3 (37%) | 2 (13%) |
| Negative | 19 (79%) | 5 (63%) | 14 (87%) |
| Prior treatment |  |  |  |
| Surgery | 18 (75%) | 6 (75%) | 12 (75%) |
| Radiation<sup>@</sup> | 23 (96%) | 8 (100%) | 15 (94%) |
| Concurrent chemoRT | 10 (42%) | 5 (63%) | 5 (31%) |
| Palliative chemotherapy | 9 (38%) | 3 (37%) | 6 (38%) |
| Disease status |  |  |  |
| Metastatic | 18 (75%) | 7 (88%) | 11 (69%) |
| Recurrent only | 6 (25%) | 1 (12%) | 5 (31%) |
| PD-L1 CPS |  |  |  |
| <1 | 16 (67%) | 5 | 11 |
| 1-19 | 3 (13%) | 0 | 3 |
| ≥ 20 | 5 (21%) | 3 | 2 |
| Anti-PD-1 Treatment |  |  |  |
| Nivolumab | 17 (71%) | 6 (75%) | 11 (69%) |
| Pembrolizumab* | 3 (13%) | 1 (12.5%) | 2 (13%) |
| PD-1 + chemotherapy* | 2 (8%) | 1 (12.5%) | 1 (6%) |
| PD-1 + lenvatinib<sup>^</sup> | 1 (4%) | 0 | 1 (6%) |
| PD-1 + GSK3359609<sup>#</sup> | 1 (4%) | 0 | 1 (6%) |

Data are median (range) or n (%). RT= radiotherapy. PD-L1= programmed death-ligand 1. CPS= combined proportion score. <sup>@</sup>Including patient who received concurrent chemoRT \*KEYNOTE-048 (NCT02358031) <sup>^</sup>LEAP-010 (NCT04199104), <sup>#</sup>INDUCE-1 (NCT02723955).

Treatment regimens included single agent nivolumab (n=17), single agent pembrolizumab (n=3) and combination anti-PD-1 therapy plus chemotherapy, lenvatinib or GSK3359609 (n=4). Prior radiation and systemic therapy is highlighted in supplementary figure 1. Majority of patients had distant metastatic disease prior to commencement of anti-PD-1 based therapy (n=18, 75%) with the lung being the most common site of metastases and 6/24 (25%) patients had locally recurrent disease only (Supplementary Figure 1). Using our response criteria, 8/24 (33%) patients were categorised as responders: one patient with CR, three patients with PR and 4 patients with SD >6 months. Of the 16/24 (67%) non-responders, one patient had SD <6 months, 11 had PD and four patients had clear clinical disease progression prior to progress imaging (RECIST not evaluable). Of the three patients who had paired primary and metastases samples, one patient was a responder (Patient ID 14), and two patients were non-responders (Patient ID 02 and 15).

## The cellular landscape of HNmSCC remain similar across different aetiologies and sites

HNmSCC is associated with different aetiologies, mainly smoking and Human Papilloma Virus (HPV) that can potentially impact the cellular landscape of the tumour environment. We first performed a comparative analysis of the overall cellular landscape according to tumor site and p16 status. Manual gating of the populations based on their phenotype showed tumor cells and immune cells to be the major constituents of the TME (Figure 1B-D, Supplementary Figure 2-3). Amongst immune populations CD4+ T cells and macrophages were the dominant immune cell populations comprising of 31.1% and 20.3% of all CD45+ cells, respectively (Figure 1C, Supplementary Figure 2-3). Interestingly, there were twice as many CD4+ T cells compared to the numbers of CD8+ T cells. Importantly, the overall TME landscape remained similar across sites (Figure 1B-D, Supplementary Figure 2-3) and p16 status (Figure 1B-D, Supplementary Figure 2-3). There were no significant differences in CD8+, CD4+ T cells, macrophages, monocytes or DCs between different aetiologies. However, there was a trend towards increased CD4+ and CD8+ T cells in p16 positive oropharyngeal SCC (n=5) and increased CD4-CD8-T cells in p16 negative tumors (n=19, Figure 1C, Supplementary Figure 2-3). There was also a trend towards increased B cells and malignant squamous cells in laryngeal tumors (n=3, Figure 1B-D, Supplementary Figure 2-3).

**Figure 1.**
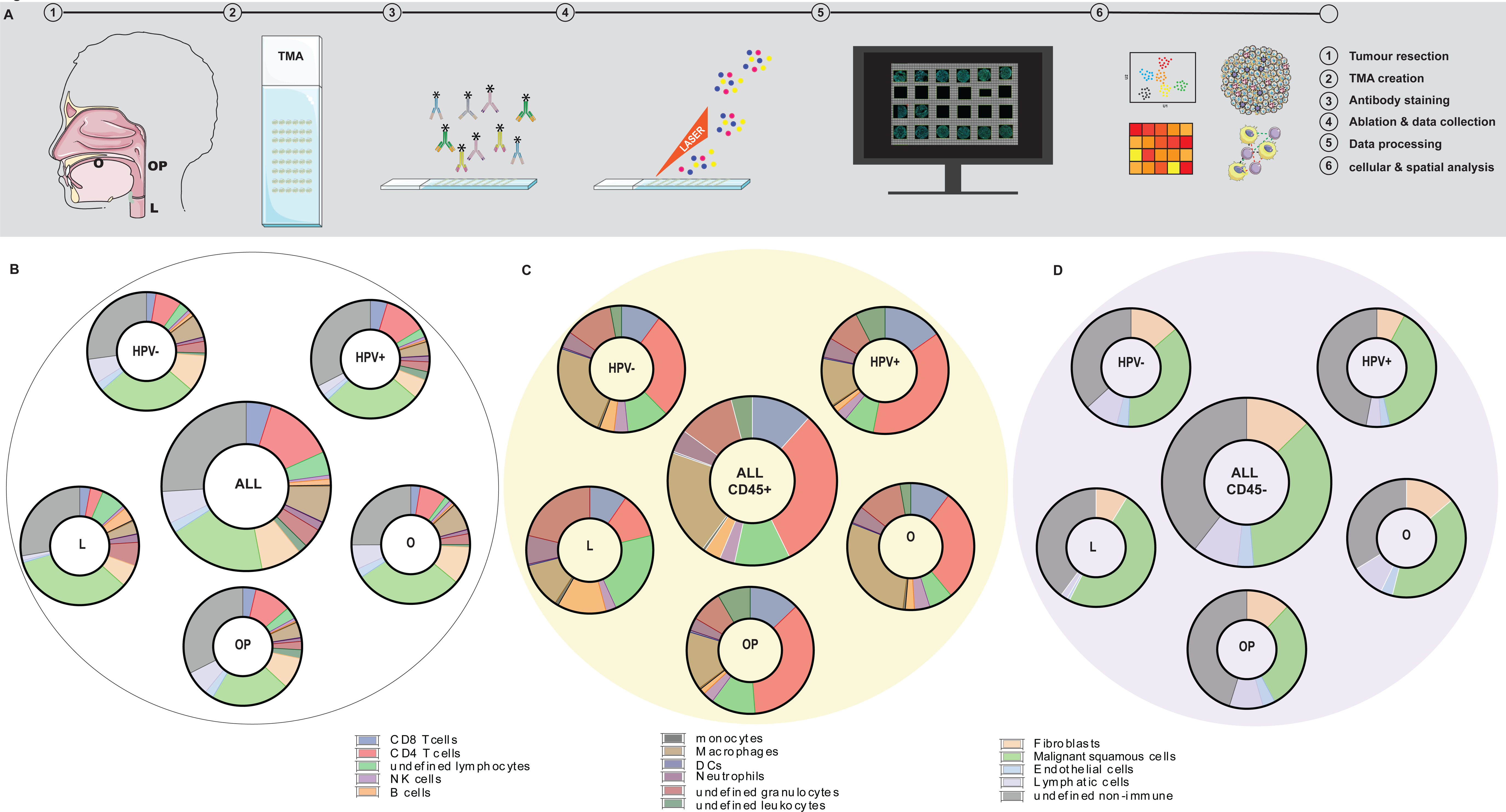
An experimental pipeline to assess cell infiltration and create a cellular map of the Head and Neck mucosal squamous cell carcinoma TME. **A** (1) 24 HNmSCC patients with Oral (O), Oropharyngeal (OP) or Laryngeal (L) tumors underwent resection. (2) 1mm^2^ intra-tumoral cores were selected from each patient in duplicate to create a tissue microarray. (3) Sections were stained with antibodies conjugated to heavy metal tags which were then (4) ablated by a Hyperion imaging mass cytometer and the data collected. (5) Data was processed using .mcd viewer and (6) cellular and spatial analysis was performed using cell profiler, FlowJo, R studio. Summary of **B** ALL cells, **C** CD45+ cells (CD8+ T cells, CD4+ Tcells, undefined lymphocytes, NK cells, B cells, monocytes, macrophages, DCs, neutrophils, undefined granulocytes, undefined leukocytes) and **D** CD45-cells (Fibroblasts, malignant squamous cells, endothelial cells, lymphatic cells, undefined non-immune) in the TME of all HNmSCC studied and grouped according to HPV status, HPV+ n=5, HPV-n=19, and location of tumor, O n=14, OP n=10 or L n=3.

## Central memory T cells and B cells are significantly increased in tumors of responders compared to non-responders

Various compositions of immune cells in tumours at baseline have been associated with response to checkpoint blockade therapies in other cancers (7). We therefore next determined both the immune and non-immune cellular composition of tumours pre-anti-PD1 treatment, between patients who subsequently responded to anti-PD-1 treatment (n=9 tumors) and those who did not (n=18 tumors). There were no significant differences in the overall density and proportion of CD45+ immune cells (Figure 2A) including CD8+ T cells (Figure 2C), regulatory T cells (Figure 2D) and innate immune cell populations, including monocytes (Figure 2F), NK cells (Figure 2G), neutrophils (Figure 2H), dendritic cells (Figure 2I) and macrophages (Figure 2J) between responders and non-responders.

**Figure 2.**
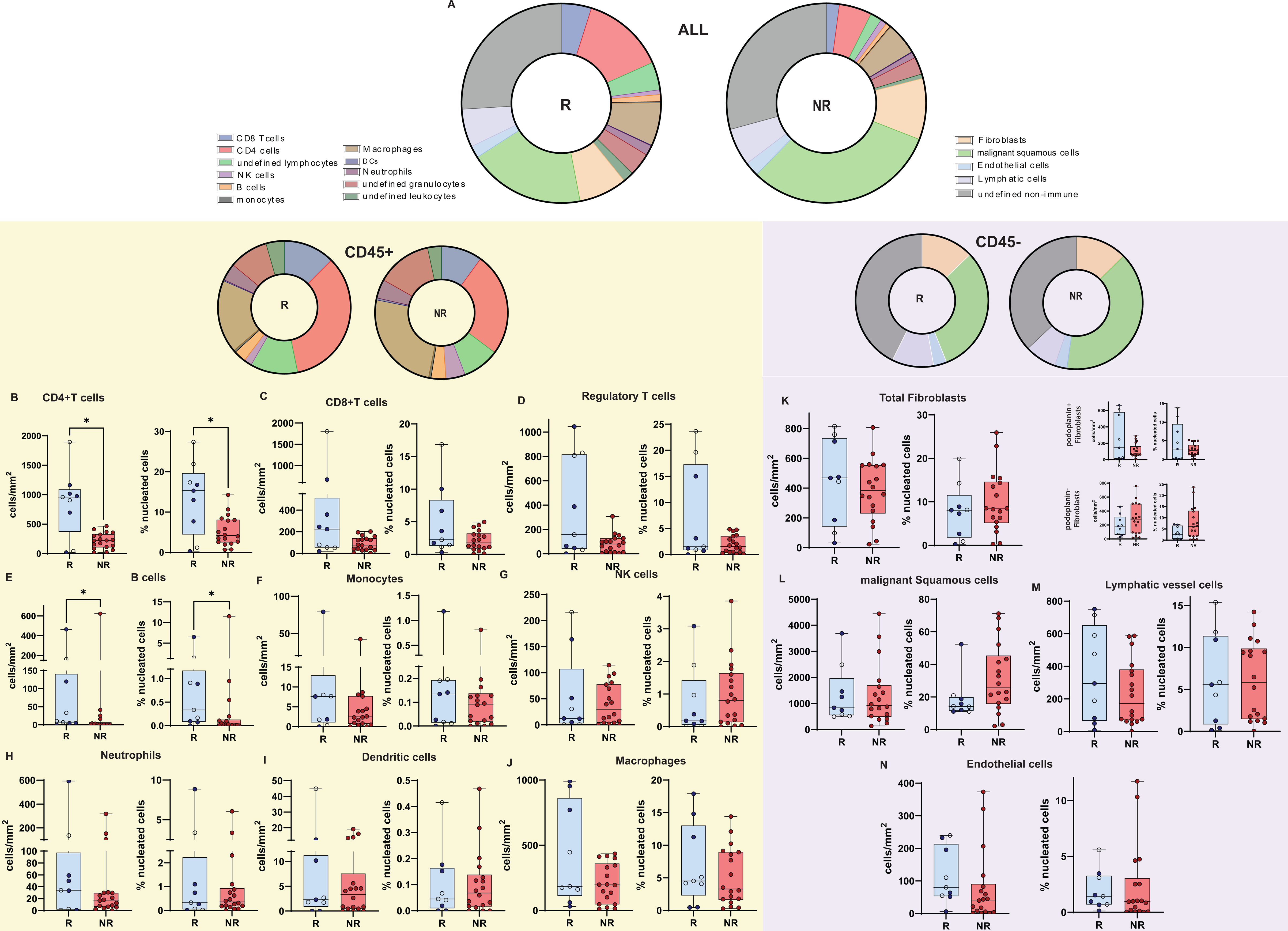
The cellular composition of the HNmSCC TME in responders and non-responders to anti-PD1 therapies. **A** Summary of ALL cells, CD45+ cells (CD8+ T cells, CD4+ T cells, undefined lymphocytes, NK cells, B cells, monocytes, macrophages, DCs, neutrophils, undefined granulocytes, undefined leukocytes) and CD45-cells (Fibroblasts, malignant squamous cells, endothelial cells, lymphatic cells, undefined non-immune) in the TME grouped according to response to anti-PD1 therapy, Responders (R) n=9 tumors, non-responders (NR) n=18 tumors. Density and proportions of immune cells according to response to anti-PD1 therapy, R (stable disease represented by open circles ○, complete and partial response by closed circles ●), NR including **B** CD4+ T cells, **C** CD8+ T cells, **D** Regulatory T cells, **E** B cells, **F** Total monocytes, **G** NK cells**, H** Neutrophils**, I** Dendritic Cells**, J** Macrophages. Density and proportions of non-immune cells according to response to anti-PD1 therapy, R, NR including **K** Total Fibroblasts, podoplanin+ and podoplanin-Fibroblasts, **L** malignant squamous cells, **M** Lymphatic vessel cells and **N** Endothelial cells. Mann-Whitney statistical test was applied to determine significance *p<0.05.

However, there was a significant increase in the density and proportion of CD4+ T cells (Proportion (Prop): p=0.014, Density (Dens): p=0.041, Figure 2B), B cells (Prop: p=0.020, Dens: p=0.036, Figure 2E) between responders compared to non-responders. Closer examination of CD4+ T cells revealed significant increases in the density of CD4+ Central memory (T_CM_) (p=0.046) and effector-memory (T_EM_) (p=0.035) T cells in responders (supplementary Figure 4C). However, no significant differences were found between responders and non-responders when comparing other CD4+ T cell subsets analysed including terminally-differentiated effector memory (T_EMRA_), naïve and tissue-resident T cells (supplementary Figure 4C). Examination of CD8+ T cell subsets revealed there was a significant increase in the proportion and density of CD8+ T_CM_ T cells (p=0.030, p=0.022) in responders when compared to non-responders (supplementary Figure 4B). However, no significant differences were found when comparing other CD8+ T cell subsets, T_EMRA_, naïve, T_EM_ and tissue-resident (supplementary Figure 4B) T cells.

The overall composition of non-immune cells across all HNmSCC patient subgroups was also mapped as shown above (Figure 2A). There was no difference in proportions and densities of total fibroblasts (panCK-p40-CD45-FXIIIa+) and podoplanin+ or podoplanin-fibroblasts (Figure 2K), malignant squamous cells (CD45-, positive for either or both panCK and p40) (Figure 2L), lymphatic cells (panCK-p40-CD45-CD31+podoplanin+) (Figure 2M) and endothelial cells (panCK-p40-CD45-CD31+podoplanin-) (Figure 2N) between responders and non-responders. Taken together here we demonstrate that pre-treatment immune cell composition, not non-haematopoietic cell composition of tumours could play a critical role in response to anti-PD-1 treatment in HNmSCC.

## Pre-treatment PD-1+ and PD-L1+ T and B cell numbers are associated with response to anti-PD-1

Accurately quantifying the immune cell populations reacting to tumour cells has been a major challenge. However, the expression of PD-1 on T cells has been associated with tumour reactivity in some cancers (8). Further categorisation of T cells on the basis of PD-1 expression revealed that the number of PD-1+ Total CD8+ T cells (Prop: p=0.048, Dens: p=0.017) as well as PD-1+ CD8+ T_CM_ (Prop: p=0.0027, Dens: p=0.001), CD8+ T_EM_ (Prop: p=0.0037, Dens: p=0.0035) and tissue-resident CD8+ T cells (Dens: p=0.045) was significantly higher in responders when compared to non-responders (Figure 3A).

**Figure 3.**
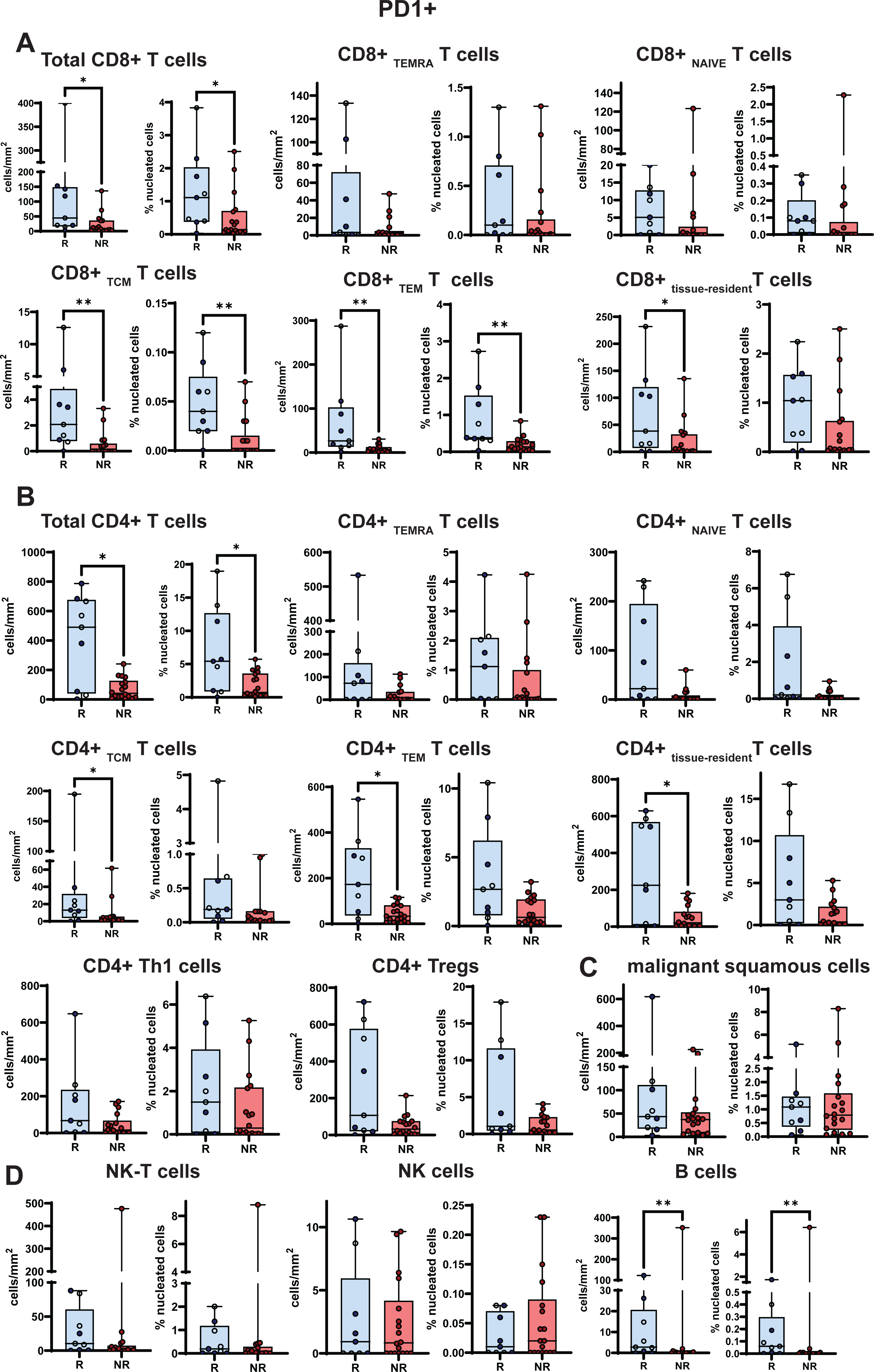
Specific PD-1+ immune cell subsets are present in the TME of HNmSCC patients that respond to anti-PD1 therapies. Density and proportions of specific PD-1+ immune cells according to response to anti-PD1 therapy, R (stable disease= open circles ○, complete and partial response= closed circles ●), NR including **A** PD-1+ Total CD8+ T cells and PD-1+ CD8+ _TEMRA_, CD8+ _NAIVE_, CD8+ _TCM_, CD8+ _TEM_, CD8+ _Tissue-resident_ T cells, **B** PD-1+ Total CD4+ T cells and PD-1+ CD4+ _TEMRA_, CD4+ _NAIVE_, CD4+ _TCM_, CD4+ _TEM_, CD4+ _Tissue-resident,_ CD4+ Th1 T cells and CD4+ Tregs, **C** PD-1+ malignant squamous cells, **D** PD-1+ NK-T cells, NK cells and B cells. Mann-Whitney statistical test was applied to determine significance. *p<0.05, **p<0.01.

PD-1+ total CD4+ T cells (Prop: p=0.026, Dens: p=0.020) as well as PD-1+ CD4+ T_CM_ (Dens: p=0.032), CD4+ T_EM_ (Dens: p=0.020) and tissue-resident CD4+ T cells (density: p=0.041) was also significantly higher in responders when compared to non-responders (Figure 3B).

In light of the increased number of PD-1+ T cell subsets, we also analysed PD-1+ in the other immune cells. There was a significant increase in the proportion and density of PD-1+ B cells (Prop: p=0.004, Dens: p=0.004) in responders compared to non-responders (Figure 3D) but no difference was found in NK or NK-T cells (Figure 3D). PD-1+ malignant squamous cells were present in similar proportions in both responders (mean 1.31% nucleated cells) and non-responders (mean 1.28% nucleated cells) (Figure 3C).

We then compared the differential expression of PD-L1 within the TME. PD-L1+ CD8+ T_CM_ (Prop: p=0.006, Dens: p=0.006) and PD-L1+ CD8+ T_EM_ (Prop: p=0.013, Dens: p=0.007) were significantly higher in responders when compared to non-responders but no difference was observed in the number of PD-L1+ total CD8+ T cells or PD-L1+ CD8+ _TEMRA,_ CD8+ _NAÏVE_ or CD8+ _Tissue Resident_ T cells (Supplementary Figure 5A). PD-L1+ total CD4+ (density: p=0.027) and CD4+ T_EM_ (Dens: p=0.023) T cells were significantly higher in responders when compared to non-responders (Supplementary Figure 5B). A trend was observed for increase in PD-L1+ CD4+ T_EM_ cells and CD4+ Tregs, however, this was not statistically significant (Supplementary Figure 5B). There was also a significant increase in the proportion and density of PD-L1+ B cells (Prop: p=0.009, Dens: p=0.005) in responders compared to non-responders but no difference was found in the number of PD-L1+ malignant squamous cells, NK, NK-T cells, macrophages or Dendritic cells (Supplementary Figure 5C-D).

## T cells interact with phenotypically distinct HNmSCC cells in responders but spatially avoid these cells in non-responders

It has recently become clear that in addition to the type of immune cell and the number of these cells in the tissue, their spatial interactions could also be critical in determining the outcomes of immune modulatory therapies (9). We therefore performed a spatial mapping of cellular interactions. Unbiased clustering analysis identified 12 distinct clusters of immune and non-immune cells within the TME (Figure 4A). Based on their phenotype (Figure 4B), cell populations were annotated to CD8+ T cells, CD4+ T cells, CD4+ Tregs, B cells, and neutrophils as the major immune cells within the TME. In addition, clustering also identified endothelial cells (PanCK-CD45-CD31+), fibroblasts (PanCK-CD45-FXIIIA+) and four phenotypically distinct malignant squamous cell populations (Figure 4A-B). These phenotypically distinct malignant squamous cell populations include: malignant squamous cell-1 (p40+Lag3+CD44+OX40+ICOS+IDO+PDL1+Ki67+), malignant squamous cell-2 (p40+Lag3+ICOS+OX40+CD103+DC-SIGN+PDL1+Ki67+), malignant squamous cell-3 (p40+PDL1+), and malignant squamous cell-4 (p40+Lag3+CD44+ CADM1+IDO+CD103+PDL1+Ki67+) (Figure 4B).

**Figure 4.**
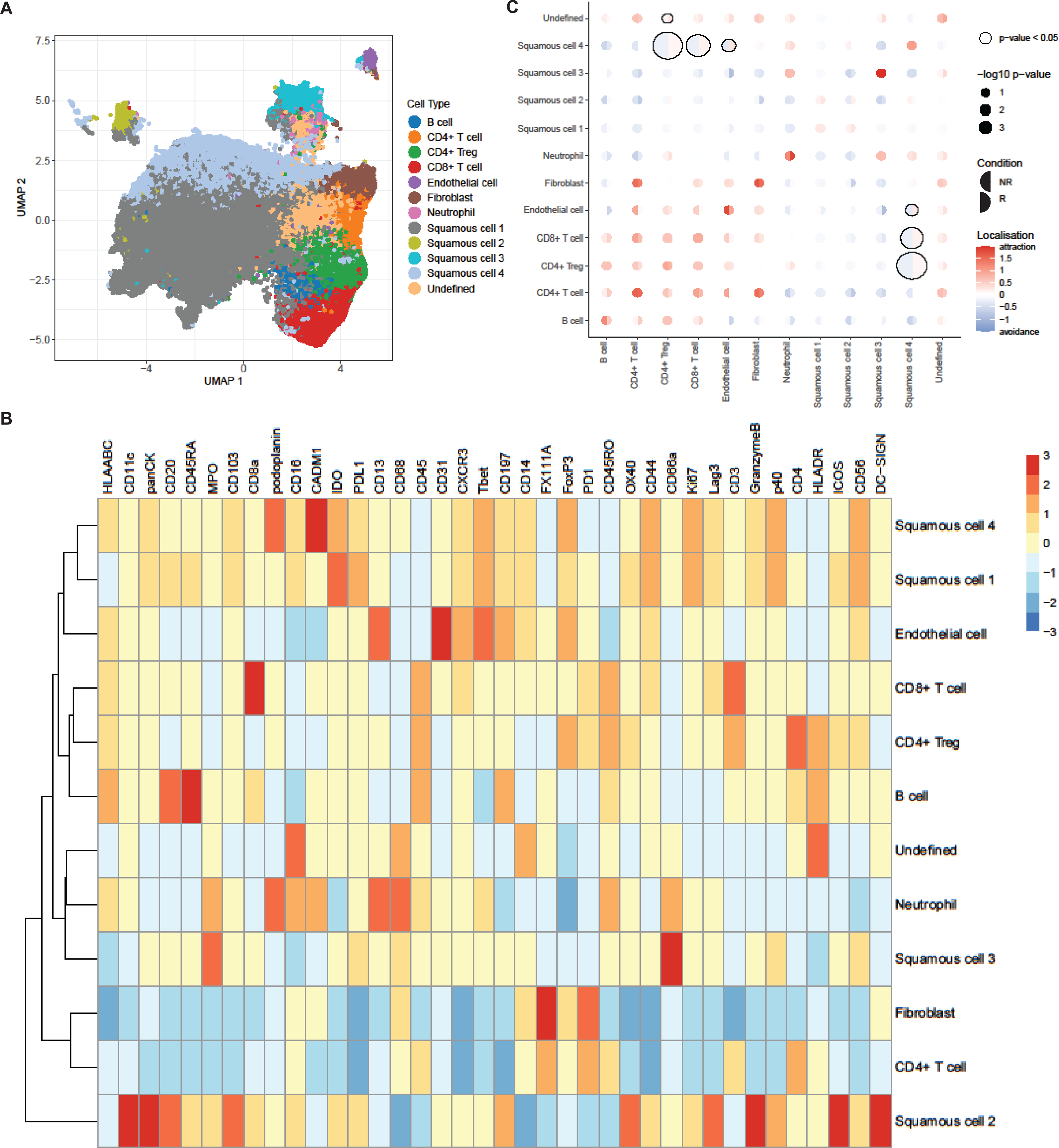
Spatial Analysis reveals significant cell-interactions in the immune landscape of Responders to anti-PD1 ICI therapy in HNmSCC. **A** Representation of cell clusters identified using UMAP analysis of tumor samples from R (n=9 tumors), NR (n=18 tumors) with clusters indicated by colour visualised in **B** heatmap of mean marker expression for each K-means cluster. Cells are phenotyped by their marker expression as B cell, CD4+ T cell, CD4+ Treg, CD8+ T cell, Endothelial cell, Fibroblast, Neutrophil, malignant squamous cell-1, malignant squamous cell, malignant squamous cell-3, malignant squamous cell-4 and undefined. **C** Cell interactions (red) or avoidance (blue) in non-responding, NR ◖ and responding, R ◗ HNmSCC samples as identified by spicyR package (Bioconductor), ○ p value <0.05, circle size indicates magnitude of statistical significance.

Geographically mapping individual cell clusters revealed clear interactions between some cells and clear avoidance between others (Figure 4C). Comparing the interactions and avoidance between cells in responders and non-responders revealed three distinct interactions. For example, malignant squamous cell-4 (p40+Lag3+CD44+ CADM1+IDO+CD103+PDL1+Ki67+) were significantly interacting with CD8+ T cells (CCR7-CD45RA-PD-1+Lag3+Ki67+), CD4+ Tregs (PD-1+Ki67+) and an endothelial cell in responders whereas these cells were significantly avoiding each other in non-responders (Figures 4B and 4C). Interestingly, the densities of PD-1+ Tregs and endothelial cells were similar in responders versus non-responders (Figure 3B, Figure 2N), suggesting it is the T cell and endothelial cell interacting with specific phenotypically distinct subsets of malignant squamous cells predict anti-PD-1 treatment response in HNmSCC.

## Increased tertiary lymphoid-like structures are associated with response to anti-PD-1 therapy and improved survival

In addition to cellular interactions, cell clusters that commonly form tissue niches could also provide insights into how immune cells contribute to anti-PD-1 therapy response. We therefore constructed performed regional analysis of our tissue maps to understand patterns of cell aggregates present within the TME. Five distinct tissue regions were identified within the TME. To characterise these regions, the cells present in the greatest frequency in each region was determined (Figure 5A). Region-1 showed presence of malignant squamous cell-4 (p40+Lag3+CD44+CADM1+IDO+CD103+PDL1+Ki67+), a proliferating tumor cell with markers associated with aggressive biology (e.g. CD44, CADM1, Lag3, IDO) and neutrophils. Region-2 included CD8+ T cells, CD4+ T cells, CD4+ Tregs and B cells (Figure 5A) with hallmarks of TLSs. Region-3 was mainly defined by the presence of two phenotypically distinct tumor cells; proliferating (Ki67+) malignant squamous cell-1 (p40+Lag3+CD44+OX40+ICOS+IDO+PDL1+Ki67+) with adverse prognostic markers such as Lag3, CD44, OX40, ICOS, IDO and malignant squamous cell-2 (p40+Lag3+ICOS+OX40+CD103+DC-SIGN+PDL1+Ki67+), a proliferating tumor cell expressing immune response down-regulating checkpoint and markers of immune status such as Lag3, ICOS, OX40, CD103 and DC-SIGN. Region-4 had a high number of CD4+ cells, endothelial cells and fibroblasts while Region-5 had a high number of malignant squamous cell-3 (p40+PDL1+), (Figure 4D) and neutrophils.

**Figure 5.**
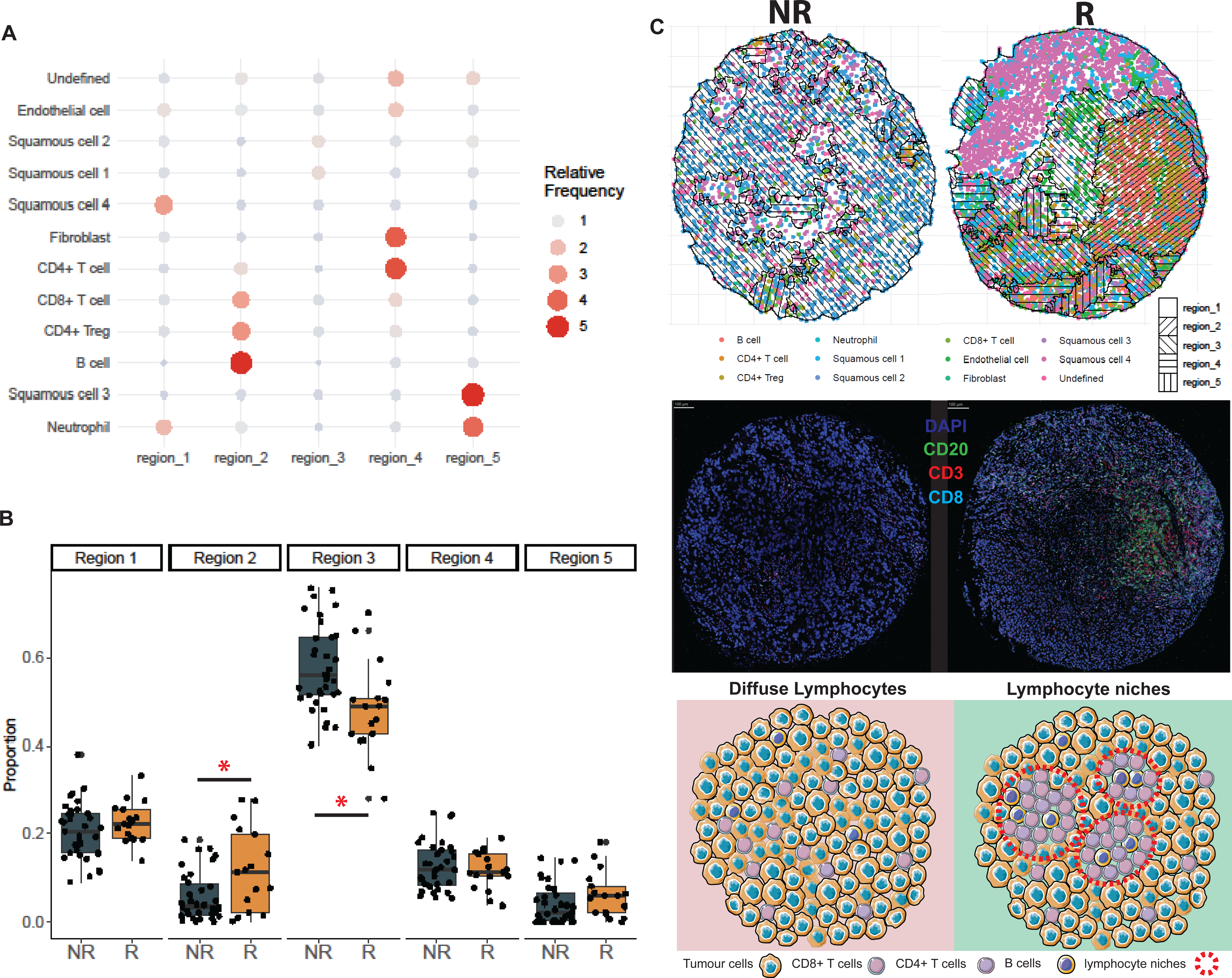
Spatial tissue-region analysis reveals tertiary lymphoid structures in the immune landscape of Responders to anti-PD1 therapy in HNmSCC. Five specific TME regions were identified within the tumor tissue of NR and R patient tumors based on magnitude and spatial placement of cell clusters. **A** Relative frequency of cells present in each region is indicated by circle size. **B** Proportion of each region present in NR and R tumor tissue. **C** Representative NR and R cell/region maps, IMC images of tissue showing DAPI= blue, CD20= green, CD3= red, CD8=cyan (scale bar 100μm) and schematic drawing demonstrating an increase of lymphocyte niches (region 2) in R compared to NR tumor tissue. Statistically significant difference *p<0.05.

We then compared the distribution of these five distinct tissue regions within the TME of responders versus non-responders. This revealed the TMEs of responders had a greater proportion of Region-2 comprising CD4+ Tregs, CD4+ and CD8+ T cells and B cells, suggestive of intratumoral TLSs (Figure 5A) and a reduced proportion of region-3 comprising a combination of multiple distinct proliferating malignant squamous cells with aggressive phenotypic features that suggest an ability to dampen anti-tumor immune responses (Figure 5B-C). The presence of TLS associated with response to anti-PD-1 therapy in the HNmSCC TME is further corroborated by topographical plots and representative images (Figure 5C) identifying TLS by the presence of immune aggregates mainly consisting of B cells, CD4+ and CD8+ T cells and CD4+ Tregs (Figure 5C). Interestingly, patients with TLS proportions >20% (n=3) were all considered exceptional responders to anti-PD1 therapy, with a PFS of 80.3 months, 26.8 months and unevaluable (unrelated death at 15.6 months).

## Discussion

This study highlights the importance of studying detailed spatial interactions within the TME and the critical limitations of quantifying cellular composition alone to predict immunotherapy response and resistance. We demonstrate that, in addition to cell densities, mapping cellular interactions and cell aggregations could also provide critical landscape information that could be used to predict response to anti-PD-1 therapy in HNmSCC. Importantly, we identify cellular interactions that are unique within the tumours of responders when compared to non-responders. We also show through regional analysis that tertiary lymphoid-like structures are significantly increased within the tumours of responders. Further to this finding, regional analysis identified other cell aggregates proportionately increased in non-responders. Together this study provides a detailed immune landscape that differentiates responders from non-responders to anti-PD-1 in HNmSCC.

TLSs are lymphocyte aggregates that develop in non-lymphoid tissues in response to antigen persistence and are associated with augmented immune-response capabilities (10). The presence and increased numbers of TLSs within tumors are associated with better prognosis in melanoma (11), lung cancer (12), clear cell renal cell carcinoma (13) and many other cancers (10). Importantly, intratumoral TLSs or their presence at tumor beds have also been found to be associated with response to immune checkpoint blockade in many cancers including melanoma (11, 14), soft tissue sarcoma (15), renal cell carcinoma and non-small cell lung carcinoma among others (12, 16). Our data demonstrates aggregation of B cells, CD4+ and CD8+ T cells, forming TLSs, in patients who responded to anti-PD-1 in HNmSCC and dispersion of B cells, CD4+ and CD8+ T cells in non-responders. Critically, in our study, HNmSCC responder patients with high proportions of TLS in their tumor (>20%) had significantly prolonged PFS. With the potential utility of CD23 in immunohistochemical staining to identify TLS there is potential for rapid translation of our findings (17). Further studies are required to determine whether these constitute early or mature TLS and the impact of TLS development state on treatment response.

Studies thus far have largely focussed on various immune mediators as potential biological predictors of response to checkpoint blockade therapies (7). Here we also provide a detailed cellular interaction and region map that is associated with response to anti-PD-1. Our data clearly demonstrates CD8+ T cells interacting with different phenotypes of tumour cells in responders, while these cells avoid each other in non-responders. One of the key factors in understanding these interactions could be the specificities of CD8+ T cells. It is possible that a higher proportion of CD8+ T cells in responders are tumour-specific, while in non-responders they are non-specific. A recent study found significant proportions of tumour infiltrating T cells to be non-specific for tumour antigens and therefore are unlikely to interact with tumour cells (18). Further supporting this possibility is our finding of increased PD-1+CD8+ T cells in responders when compared to non-responders. It is also possible TLS are able to maintain more tumor-reactive CD8+ T cells within responder tumours when compared to non-responder tumours as suggested by studies showing continuous trafficking of T cells between tumour-draining lymph nodes and tumours (19).

The density of total CD8+ T cells in tumor samples of patients receiving anti-PD1 therapy has predicted improved response and outcomes in some cancer types (20), but not all (21). Our study also did not find significant differences in the densities of total CD8+ T cells between responders and non-responders. However, a closer look at a cellular subset level, identified T_CM_ and PD-1+ CD8+ T cell subsets as significantly increased in responders, highlighting the importance of subset level analysis. It is also worth noting that CD8+ T cells expressing both TCF-1 and PD-1 have been found to be the key initiators of response to anti-PD1 therapy in both pre-clinical models and in human cancers (22, 23). Importantly, these cells were largely found within the tumour draining lymph nodes (18). Although our study did not determine TCF-1 expression, an increased density of PD-1+CD8+ T cells was found in responders and the CD8+ T cell present in the TLS region mapped in our study was also PD-1+. Besides CD8+ T cells, enrichment of tumor-reactive B cells within TLS has also been found to enhance tumor clearance in some cancers (16). Significantly increased total and PD-1+ B cells in our study also suggests a potential role for these cells in tumour control. Further studies are however required to characterise the specificities and functions of these B cells. Similarly, there is increasing evidence showing a critical role for CD4+ T cells in tumour control and in response to checkpoint blockade therapies (24). CD4+ T cells have been directly implicated in tumour killing (25, 26), providing help to CD8+ T cells, licensing antigen presenting cells and importantly facilitating tumour clearance through secretion of cytokines like IFN-γ (27). In this regard our data also suggests a potential key role for CD4+ T cells in response to anti-PD-1 in HNmSCC. Importantly, the cellular aggregation of all these cells suggests a supportive niche in which the potential initiators of response could be maintained.

Cellular composition of tumours can also inform the characteristics of the environment in which immune cells become dysfunctional and the likelihood of those tumours responding to immunotherapies. In this regard, the constituents of HNmSCCs and how much impact the aetiology or location has on immune composition has remained unknown. Generally, tumours associated with viral infections are more immunogenic and have greater immune infiltrates compared to tumours associated with lifestyle factors such as smoking (28, 29). A detailed mapping of advanced HNmSCC in this study, however, showed that the immune landscape largely remained similar irrespective of the head and neck mucosal subsite, aetiology, histopathologic features or p16 status. Our findings indicate that the aetiologically and morphologically diverse HNmSCC are similar at the level of TME. This suggests that response to anti-PD-1 in HNmSCCs could be independent of location or aetiology. This is supported by the phase III clinical trial (KEYNOTE-040), which did not demonstrate a significant difference in OS between HPV-positive (n=119, HR 0.97, 95% CI 0.63-1.49) versus HPV-negative (n=376, HR 0.77, 95% CI 0.61-0.97) subgroups(30).

While the response rate to anti-PD-1 remains low in HNmSCCs, our study has identified critical aspects of the TME that are associated with response or resistance to anti PD-1 therapy. Our spatial mapping has demonstrated significant interactions between specific cellular components within the TME, and the presence and importance of TLS in maintaining durability of response to anti-PD1 therapy. These characteristics can be used to develop biological predictors of response with greater specificity than currently available biomarkers such as PD-L1, tumor mutation burden and T-cell-inflamed gene expression profiles, in addition to strategies to overcome resistance.

## Online Methods

### Cohort selection and clinicopathologic data collection

Following human ethics approval through the Sydney Local Health District Ethics Review Committee (Ref: X19-282), patients were selected for this study based on three inclusion criteria:

1. A histopathologically proven diagnosis of recurrent or metastatic HNmSCC.
2. Availability of suitable Formalin Fixed Paraffin Embedded (FFPE) tumor specimen taken prior to commencement of anti-PD-1 treatment.
3. Treatment with either Nivolumab or Pembrolizumab (anti-PD-1) based therapies with complete clinicopathological and radiological data.

Clinicopathological data including patient demographics, prior therapy, primary site, sites or metastases, p16 status and PD-L1 (22C3) CPS was collected (Table 1 and Supplementary Figure 1) (31), (32). Anti-PD-1 therapy details including agent, duration, and treatment response were also collated (Table 1). Treatment response was determined as per RECIST 1.1, where we categorised patients who achieved complete response (CR) partial response (PR) or stable disease (SD) >6 months as responders (R) and patients who had SD <6 months and progressive disease (PD) as non-responders (NR). Patients who were unable to have progress imaging due to rapid clinical progression were also categorised as non-responders.

### Tissue processing, staining, data accrual and analysis

Haematoxylin and eosin (H&E) stained sections from FFPE tumor specimens were reviewed by a specialist anatomical pathologist. Two representative intratumoral cores were subsequently taken from each FFPE tissue block. The regions of interest sampled were the intra-tumoral advancing or infiltrating edge of the tumor and the centre of the tumor whenever possible. The core samples taken were 1mm^2^ in size. A total of 27 tumors were sampled from 24 patients, as some patients had both primary and metastatic lesions included. The 54 tissue cores were arranged into two tissue microarrays (TMA), sectioned at 7µm onto charged slides, in preparation for staining (Figure 1A).

An antibody panel was devised to capture the wide range of components of interest constituting the TME. Specific components of interest included markers of immune cells, squamous cells, non-immune cells such as stromal cells, fibroblasts and endothelial cells, immune checkpoint markers, cell proliferation and activation markers and cell signalling markers (Supplementary table 1). Validation was performed as previously published (33). Unconjugated antibodies were validated by traditional IHC on FFPE specimens to confirm accuracy. These were subsequently revalidated following heavy metal tag conjugation and concentrations adjusted. Pre-conjugated antibodies were validated, and concentrations adjusted using IMC on comparable FFPE tissue specimens prior to use on experimental samples. Stained TMAs were analysed by IMC using Hyperion^TM^ Imaging System (Standard Biotools) (Figure 1A) as previously described (33).

Data was produced as mass cytometry data files (.mcd) which were converted into .tiff files using MCDviewer (v1.0.560.6 Fluidigm). Data analysis was performed using a multi-software analysis pipeline including the open-source CellProfiler version 2.2.0 (cell segmentation) and Flowjo (licenced, Becton Dickinson & Company, version 10.8.1) (Figure 1A) as performed previously (33). We determined the proportion and densities of immune cells, tumor cells and other non-immune cells by manually gating on cells based on their positive or negative marker expression to determine each phenotype. Tumor cells were identified as positive for one or both p40 and pan-cytokeratin and non-haematopoietic (CD45-). Amongst the CD45+ immune cells, we identified all major cell types, including T cells (CD4+ and CD8+), B cell, macrophages, monocytes, dendritic cells, neutrophils, granulocytes and NK cells.

### Unbiased clustering and Spatial Analysis

Intensity values of various markers were square root transformed and normalized by trimming the mean to the 99^th^ quantile for each marker and regressing out the first principal component across all markers. Unbiased clustering was performed by applying the FuseSOM package(34) to the normalized intensity values of CD20, CD31, podoplanin, CD66, FXIIIA, CD14, CD3, CD4, CD8a, CD56, CD11c, MPO, CD45, panCK, p40 and FoxP3. This grouped similar immune, stromal, or squamous cell subgroups within the collated HNmSCC specimens. FuseSOM was run with 12 clusters and default parameters. Unbiased clustering enabled the capture of cell populations that do not fall outside of conventional pre-existing phenotypes. The presence of the cells identified through unbiased clustering was confirmed by Flowjo gating using cell surface markers present. Each cluster was visualized using Uniform Manifold Approximation and Projection for Dimension Reduction (UMAP) (16). The assigned phenotype for each cluster was based on the average expression of each marker as visualised by heatmap.

### Software and statistical analysis

Tests for spatial associations were done using various R packages. The spicyR package (17) was used to test for changes in the spatial relationships between paired combinations of cells. The lisaClust package (35) was used to identify and visualize regions of tissue where spatial associations between cell types are similar. The statistical significance of spatial interactions between the cell types in responder and non-responder groups was determined using the output from the spicyR and lisaClust R packages.

Finally, predictive statistical learning analysis was performed to predict response to therapy. The scFeatures package (36) was used to calculate the average abundance of each marker in each cluster of cells. Association of these features with response was tested using t-tests. The ClassifyR package (37) was used to predict response using the generated features.

R package (version 2.2.9) and Graphpad Prism (version 9.3.1) were used to perform statistical analysis. Mann-Whitney non-parametric tests were performed, with a statistical significance level of p<0.05 established.

## Acknowledgements

We would like to acknowledge the follow sources of funding supporting this research: The Cancer Institute New South Wales (CINSW 2020/2081), Sydney Local Health District, and the National Health and Medical Research Council.

We gratefully acknowledge Sydney Cytometry and thank the support staff in this core facility for their assistance. We gratefully acknowledge the Head and Neck Cancer Biobank (H20/113193) and thank the support staff for their assistance.

Parts of Figures. 1A and 5C were created using templates from Servier Medical Art (http://smart.servier.com/), licensed under a Creative Common Attribution 3.0 Generic License.

---

**Supplementary Figure 1.**
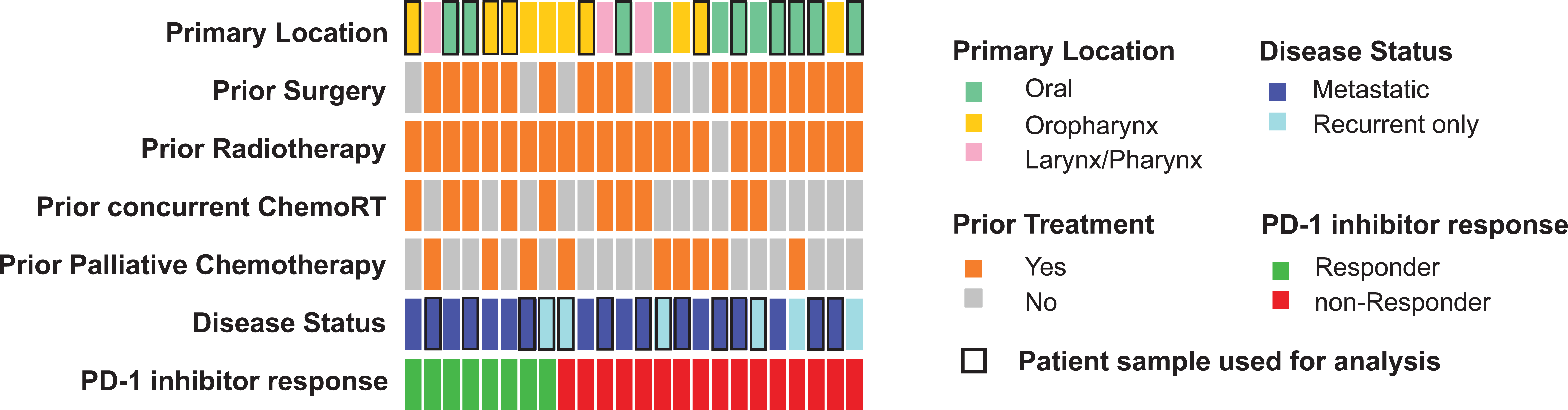
Patient summary of disease location, prior treatment, extent and subsequent anti-PD1 response. Primary location of disease (Oral, Oropharynx, Larynx/Phraynx), mode of prior treatment (Prior surgery, Radiotherapy, concurrent chemoRT, Palliative chemotherapy), extent of disease (recurrence only or metastasis), subsequent response to anti PD-1 therapy (Responder, non-Responder).

**Supplementary Figure 2.**
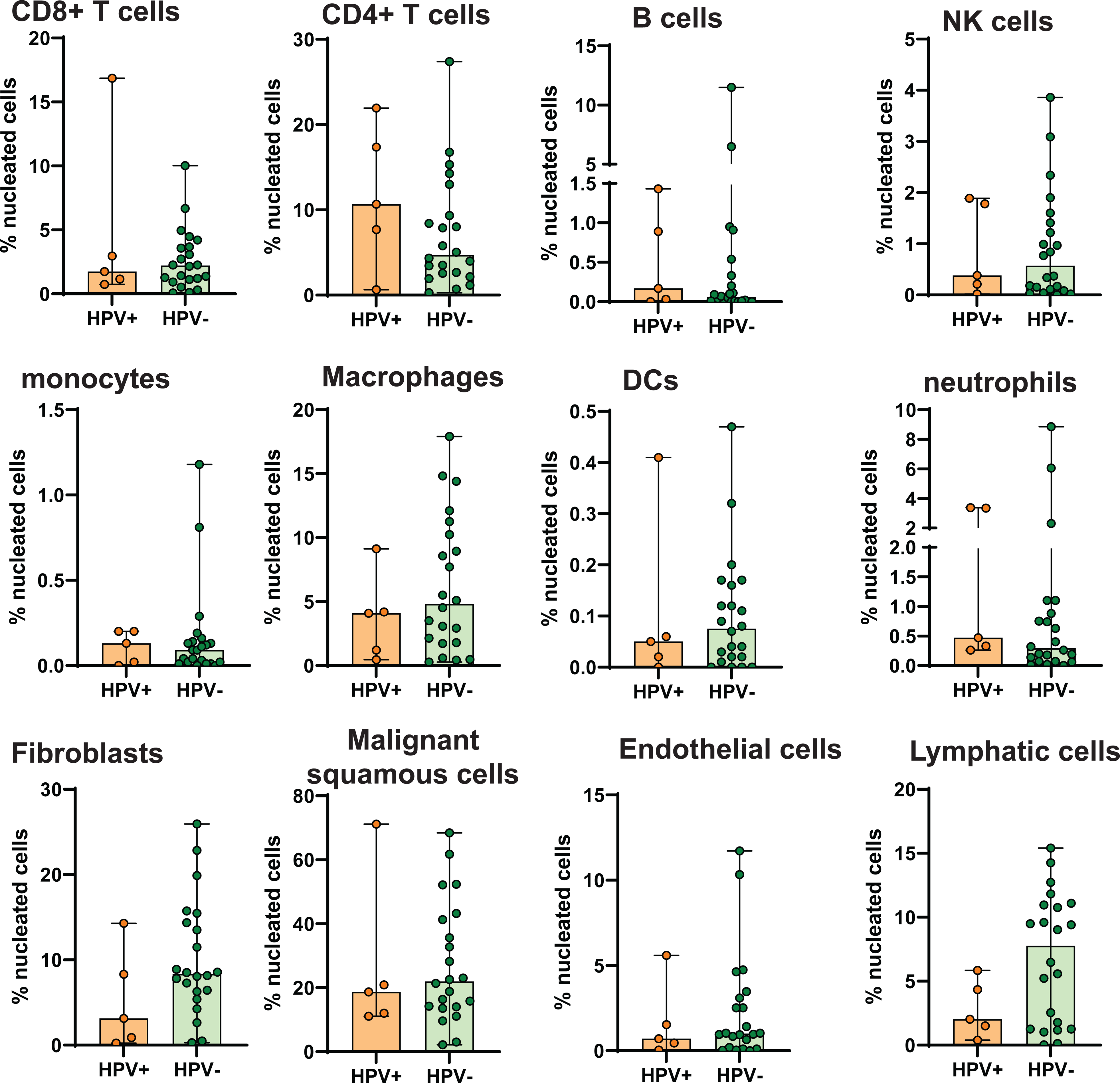
The cellular composition of the HNmSCC TME according to HPV status. Density and proportions of immune cells according to HPV status, HPV+, HPV-including CD4+ T cells, CD8+ T cells, B cells, NK cells, monocytes, Macrophages, Dendritic Cells (DCs) and neutrophils. Density and proportions of non-immune cells according to HPV (p16) status, HPV+, HPV-including Fibroblasts, malignant squamous cells, Endothelial cells and Lymphatic vessel cells.

**Supplementary Figure 3.**
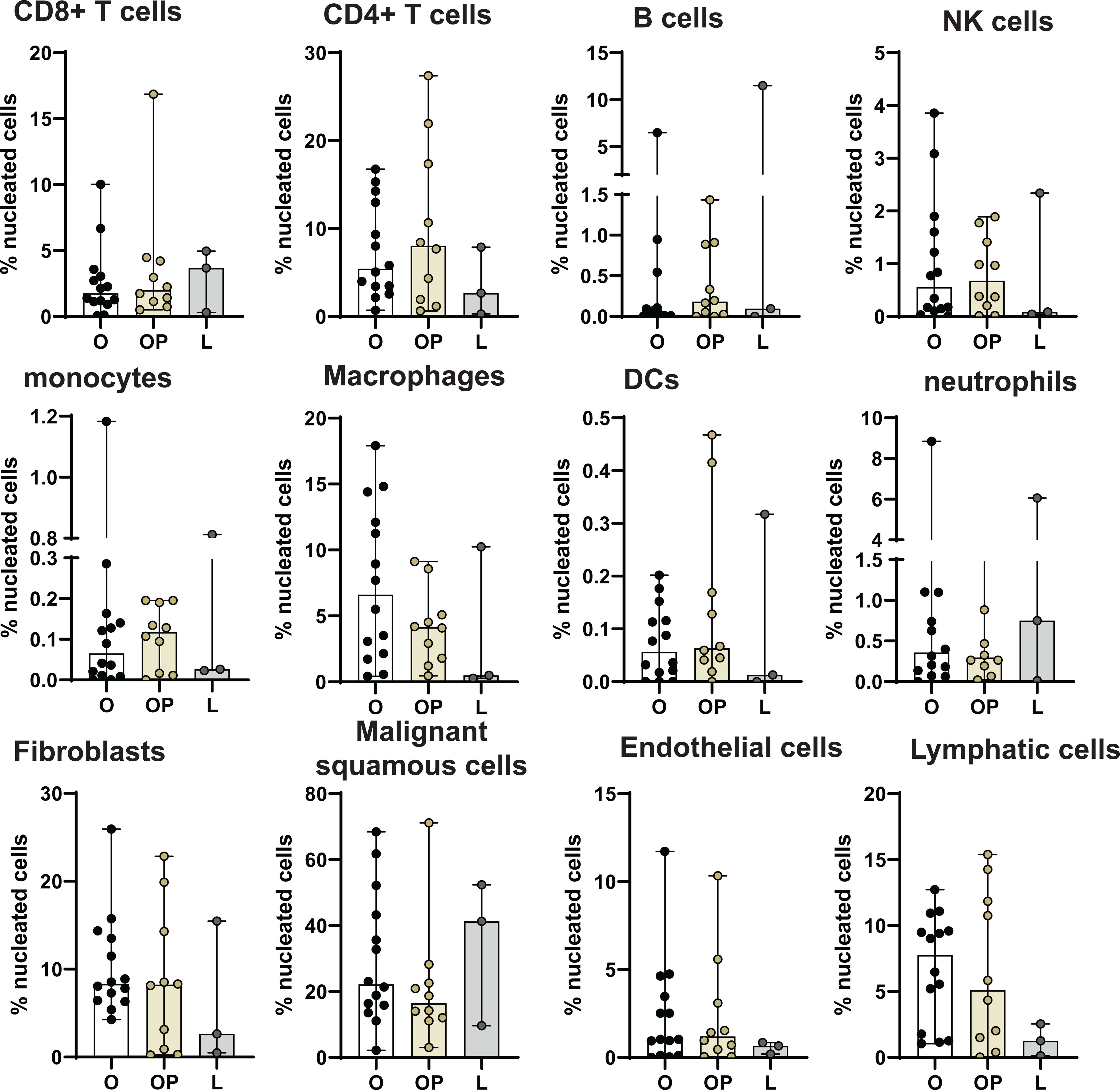
The cellular composition of the HNmSCC TME according to tumor location. Density and proportions of immune cells according to tumor location, Oral (O), Oropharyngeal (OP) or Laryngeal (L) including CD4+ T cells, CD8+ T cells, B cells, NK cells, monocytes, Macrophages, Dendritic Cells (DCs) and neutrophils. Density and proportions of non-immune cells according to tumor location, O, OP or L including Fibroblasts, malignant squamous cells, Endothelial cells and Lymphatic vessel cells.

**Supplementary Figure 4.**
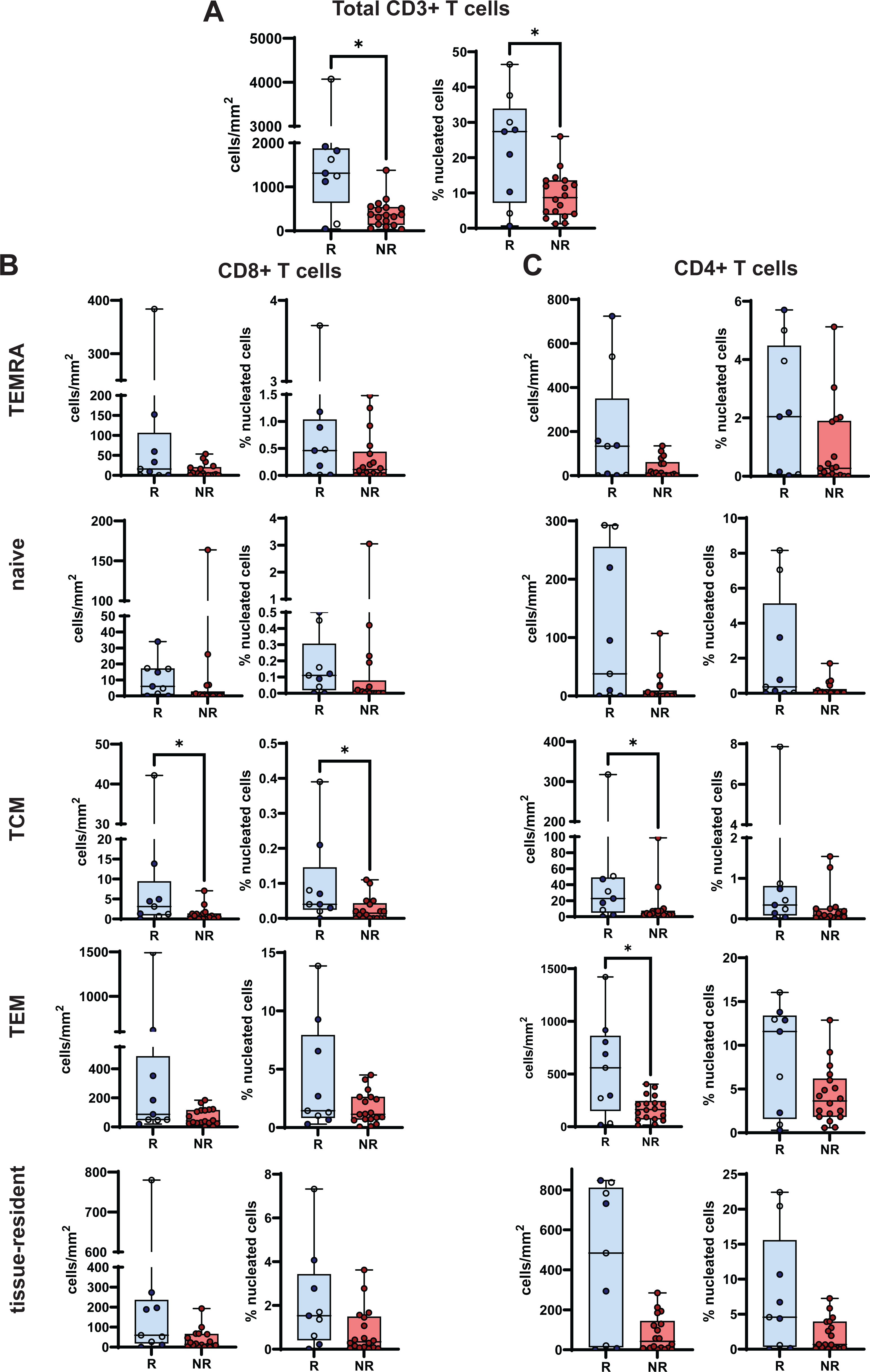
Specific T cell subsets are present in the TME of HNmSCC patients that respond to anti-PD1 therapies. Density and proportions of specific T cells according to response to anti-PD1 therapy, R (stable disease= open circles ○, complete and partial response= closed circles ●), NR, including **A** Total CD3+ T cells and **B** CD8+ _TEMRA_, CD8+ _NAIVE_, CD8+ _TCM_, CD8+ _TEM_, CD8+ _Tissue-resident_ T cells and **C** Total CD4+ T cells and CD4+ _TEMRA_, CD4+ _NAIVE_, CD4+ _TCM_, CD4+ _TEM_, CD4+ _Tissue-resident_ T cells. Mann-Whitney statistical test was applied to determine significance. *p<0.05, **p<0.01.

**Supplementary Figure 5.**
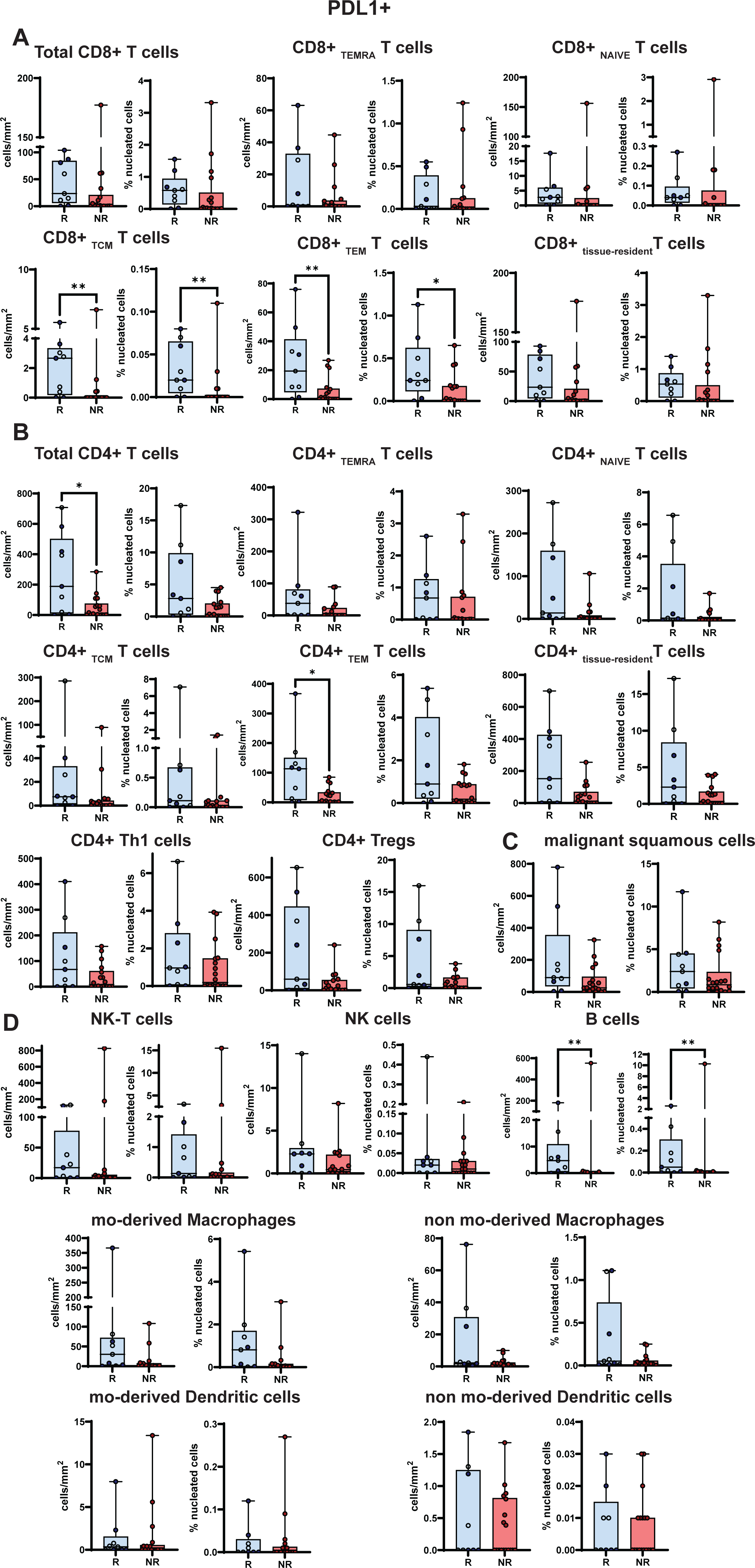
Specific PDL-1+ immune cell subsets are present in the TME of HNmSCC patients that respond to anti-PD1 therapies. Density and proportions of specific PDL-1+ immune cells according to response to anti-PDL-1 therapy, R (stable disease= open circles ○, complete and partial response= closed circles ●), NR including **A** PDL-1+ Total CD8+ T cells and PDL-1+ CD8+ _TEMRA_, CD8+ _NAIVE_, CD8+ _TCM_, CD8+ _TEM_, CD8+ _Tissue-resident_ T cells, **B** PDL-1+ Total CD4+ T cells and PDL-1+ CD4+ _TEMRA_, CD4+ _NAIVE_, CD4+ _TCM_, CD4+ _TEM_, CD4+ _Tissue-resident,_ CD4+ Th1 T cells and CD4+ Tregs, **C** PDL-1+ malignant squamous cells, **D** PDL-1+ NK-T cells, NK cells, B cells and monocyte-derived and non-monocyte-derived Macrophages and Dendritic cells. Mann-Whitney statistical test was applied to determine significance. *p<0.05, **p<0.01.

**Supplementary Table 1.** IMC antibody details.

| ***Metal Tag*** | ***Target*** | ***Antibody clone*** | ***Manufacturer*** |
| --- | --- | --- | --- |
| **113In** | HLA-ABC | EMR8-5 | abcam |
| **115In** | CD11c | D3V1E | CST |
| **139La** | Pan-cytokeratin | SPM115, SPM116 | Novus Biologicals |
| **141Pr** | CD20 | H1 | BD |
| **142Nd** | Histone H3 | Polyclonal | Novus Biologicals |
| **143Nd** | CD45RA | HI100 | Biolegend |
| **144Nd** | MPO | E1E71 | CST |
| **145Nd** | CD103 | EPR4166 | abcam |
| **146Nd** | CD8 | D8A8Y | Cell signalling |
| **147Sm** | Podoplanin | Polyclonal | RnD Systems |
| **148Nd** | CD16 | EPR16784 | abcam |
| **149Sm** | CADM1 | Poloclonal | Invitrogen |
| **150Nd** | IDO | D5J4E | CST |
| **151Eu** | PD-L1 | E13LN | CST |
| **152Sm** | CD13 | 498001 | RnD Systems |
| **153Eu** | CD68 | KP1 | Biolegend |
| **154Sm** | CD45 | D9M8I | CST |
| **155Gd** | CD31 | EPR3094 | abcam |
| **156Gd** | CXCR3 | 49801 | abcam |
| **158Gd** | T-bet | D6N8B | CST |
| **159Tb** | CD197 (CCR7) | Polyclonal | Proteintech |
| **160Gd** | CD14 | EPR3653 | abcam |
| **161Dy** | FXIIIa | Polyclonal | Affinity Biologicals |
| **162Dy** | FoxP3 | Polyclonal | RnD Systems |
| **163Dy** | PD-1 | EPR4877 (2) | abcam |
| **164Dy** | CD45RO | UCHL1 | Becton Dickinson |
| **165Ho** | OX40 | E9U70 | CST |
| **166Er** | CD44 | BJ18 | Biolegend |
| **167Er** | CD66a | YTH71.3 | abcam |
| **168Er** | Ki67 | Polyclonal | RnD Systems |
| **169Tm** | LAG3 | D2G40 | CST |
| **170Er** | CD3 | Polyclonal | abcam |
| **171Yb** | Granzyme B | EPR8260 | abcam |
| **172Yb** | p40 | BC28 | Biocare |
| **173Yb** | CD4 | EPR6855 | abcam |
| **174Yb** | HLA-DR | EPR3692 | abcam |
| **175Lu** | ICOS | D1K2T | CST |
| **176Yb** | CD56 | EPR2566 | abcam |
| **209Bi** | DC SIGN | polyclonal | abcam |
| **191Ir** | DNA intercalator | - | Fluidigm |
| **193Ir** | DNA intercalator | - | Fluidigm |

## Notes

Conflicts of interest: JHL declares Honorarium from MSD, BMS, AstraZeneca, Conference support from Novartis, Advisory Board MSD, Sanofi.

